# Developmental cell death of cortical projection neurons is controlled by a Bcl11a/Bcl6-dependent pathway

**DOI:** 10.1101/2021.10.06.463323

**Authors:** C. Wiegreffe, T. Wahl, N. S. Joos, J. Bonnefont, P. Vanderhaeghen, P. Liu, S. Britsch

## Abstract

Developmental neuron death plays a pivotal role in refining organization and wiring during neocortex formation. Aberrant regulation of this process results in neurodevelopmental disorders including impaired learning and memory. Underlying molecular pathways are incompletely determined. Loss of Bcl11a in cortical projection neurons induces pronounced cell death in upper-layer cortical projection neurons during postnatal corticogenesis. We used this genetic model to explore genetic mechanisms by which developmental neuron death is controlled. Unexpectedly, we found Bcl6, previously shown to be involved in transition of cortical neurons from progenitor to postmitotic differentiation state to provide a major check point regulating neuron survival during late cortical development. We show that Bcl11a is a direct transcriptional regulator of *Bcl6*. Deletion of *Bcl6* exerts death of cortical projection neurons. In turn, reintroduction of *Bcl6* into *Bcl11a* mutants prevents induction of cell death in these neurons. Together, our data identify a novel Bcl11a/Bcl6-dependent molecular pathway in regulation of developmental cell death during corticogenesis.

## Introduction

Developmental cell death (DCD) occurs in all animals and organs. It is part of a homeostatic balance between generation and elimination of cells. DCD provides a major check point for quality control allowing selective removal of either defective, mis-integrated or no longer required cells (Causeret et al., 2018; Wong and Marin, 2019). During development of the mammalian neocortex excess numbers of neurons are generated. Supernumerary neurons are eliminated during two distinct waves of apoptosis. In mice, a first wave of DCD occurs around E14 and affects predominantly proliferating neuron precursors (Blaschke et al., 1996; de la Rosa and de Pablo, 2000; Roth et al., 2000). During a second wave, corresponding to the first two postnatal weeks in rodents, approximately 30% of postmitotic cortical neurons are eliminated by DCD (Southwell et al., 2012; Verney et al., 2000). Within this period, entire neuron populations, as for example Cajal-Retzius cells, which transiently serve as signaling centers, are removed by DCD (Chowdhury et al., 2010; Ledonne et al., 2016), while in other neuron types, like cortical projection neurons (CPN), DCD adjusts definitive neuron numbers and refines immature synaptic circuits (Blanquie et al., 2017; Wong et al., 2018). In the neocortex dysregulated DCD has been shown to be associated with a wide spectrum of neurodevelopmental disorders, including major structural changes as well as structurally more subtle defects, like autism-spectrum disorders and intellectual disability (Eriksson et al., 2001; Kuida et al., 1996; Nakamura et al., 2016; Wei et al., 2014). DCD acts cell-type specific and is spatio-temporarily highly restricted suggesting complex molecular regulation. In contrast to the peripheral nervous system, where target-derived neurotrophic signals have been extensively demonstrated to play a key role in regulation of neuron survival (Huang and Reichardt, 2001), the molecular controls of DCD within the central nervous system (CNS) are incompletely determined. Electrical and synaptic activity have been shown to confer survival signals onto postmitotic cortical neurons (Blanquie et al., 2017; Denaxa et al., 2018; Priya et al., 2018; Wong et al., 2018). Transcription factor cascades as well as secreted signaling molecules are of key importance for the development of the neocortex. It is, however, unclear, how these regulatory networks are connected to DCD.

*Bcl11a* (*Ctip1*) encodes a zinc-finger protein that regulates transcription through interaction with COUP-TF proteins as well as direct, sequence-dependent DNA binding (Avram et al., 2002). We recently demonstrated that postmitotic upper-layer CPN require expression of Bcl11a for early postnatal survival. Cre/loxP dependent ablation of *Bcl11a* in CPN results in massive increase of apoptosis between P4 and P6 selectively in upper-layer CPN (Wiegreffe et al., 2015).

In this study we employed *Bcl11a* mutation in CPN as highly selective genetic tool to systematically identify downstream-candidate genes involved in the regulation of DCD in postmitotic CPN. Using comparative transcriptome analyses we found that Bcl6, previously reported to be involved in transition of cortical neurons from progenitor to postmitotic differentiation state (Bonnefont et al., 2019; Tiberi et al., 2012), is downregulated in *Bcl11a* mutant upper-layer CPN. Furthermore, we show Bcl11a to directly bind to a conserved promotor element of the *Bcl6* gene. Knock-out of *Bcl6* in postmitotic CPN induces their apoptosis. In turn, reintroduction of *Bcl6* into *Bcl11a* mutant CPN prevents these neurons from apoptosis. Finally, we show *Foxo1* to be downregulated in *Bcl6* mutant CPN. The Foxo transcription factor family is well established for their ability to regulate apoptosis in neurons (Santo and Paik, 2018) suggesting Bcl6 to regulate DCD at least in part through Foxo1 function. Taken together, in this study we demonstrate that DCD of postmitotic upper-layer CPN is controlled by a novel Bcl11a/Bcl6-dependent transcriptional pathway.

## Results

### Identification of downstream candidate targets of Bcl11a

We used *Bcl11a^F/F^; Emx1^IRESCre^* brains as a model to identify genes that play a role in postnatal survival of projection neurons in the somatosensory neocortex. *Bcl11a* mutant brains display robust increase of apoptosis during the second wave of DCD in upper cortical layers at postnatal stages (Wiegreffe et al., 2015). Using laser capture microdissection we specifically isolated cortical layers 2-4 of *Bcl11a* mutant and control brains at P2 (Fig. 1A, B), a stage when the second wave of apoptosis has not yet been initiated (Blanquie et al., 2017) and cell death is not yet increased in *Bcl11a* mutants (Wiegreffe et al., 2015). We then performed a differential expression analysis using microarrays and identified a set of 137 differentially expressed (DE) genes that were subjected to a GO overrepresentation test, which revealed genes involved in axon guidance, cell-cell adhesion, and regulation of cell communication (Fig. 1C; Fig. S1; Fig. S2A; Tab. S1). To verify the validity of the experimental approach, selected candidate genes were tested by quantitative real-time PCR and RNA *in situ* hybridization. *Cdh6*, *Cdh12*, *Efna5*, and *Pcdh9* that were identified as downregulated were verified by this approach (Fig. S2B, C). In addition, *Cdh13*, *Flrt2*, *Flrt3*, and *Slit2* were verified as upregulated (Fig. S2B, D). Together, these results show that our genetic approach consistently identified DE genes in upper cortical layers of *Bcl11a* mutant brains that could directly or indirectly be involved in the regulation of developmental apoptosis.

**Figure 1.**
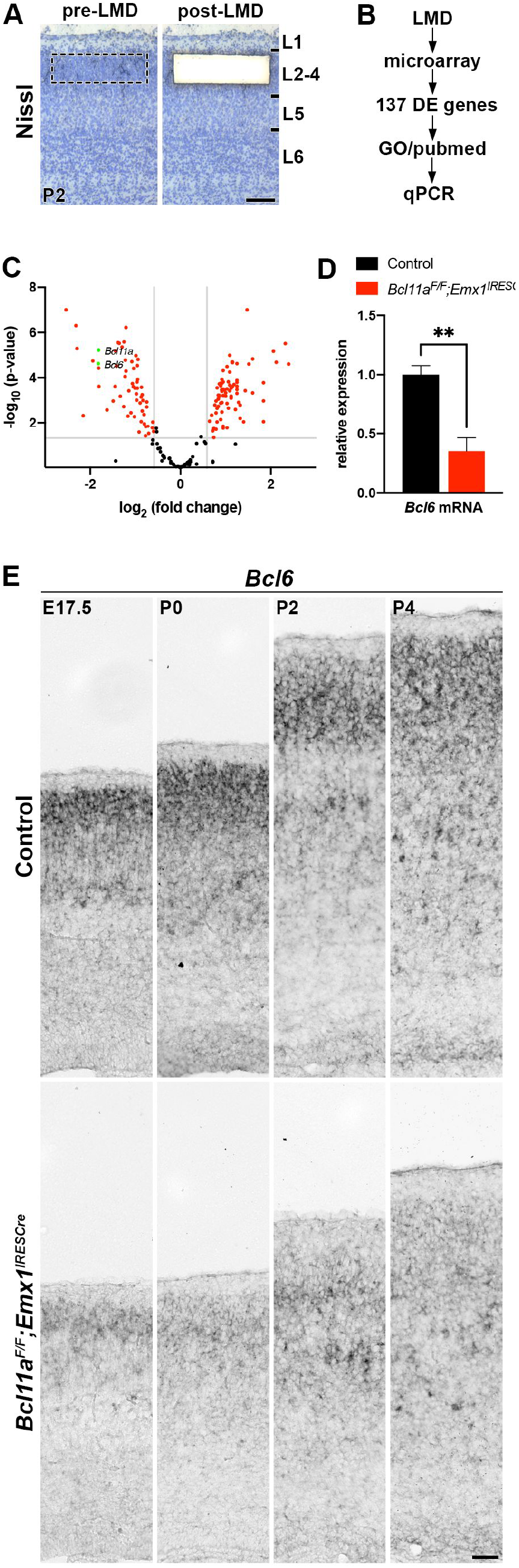
Identification of downstream candidate target genes of Bcl11a in superficial cortical layers at early postnatal development. (A) Cortical layers 2-4 was isolated by laser microdissection in 4 replicates from *Bcl11a^F/F^;Emx1^IRESCre^* and control neocortex. (B) Gene expression was compared using microarrays. From a set of 139 differentially expressed (DE) genes candidate targets were selected based on gene ontology (GO) and Pubmed analyses and verified by quantitative real-time PCR and RNA *in situ* hybridization. (C) Volcano plot showing DE genes (red). Those not significantly changed (fold change < 1.5; *p* > 0.05) are shaded black. Bcl11a and Bcl6 are highlighted in green. (D) Relative *Bcl6* mRNA expression level determined by quantitative real-time PCR is decreased in laser microdissected cortical tissue of P2 *Bcl11a^F/F^;Emx1^IRESCre^* compared to control brains (n = 4). (E) RNA *in situ* hybridization showing downregulation of *Bcl6* expression in *Bcl11a^F/F^;Emx1^IRESCre^* compared to control neocortex at E17.5, P0, P2, and P4. Student’s t test; ** p > 0.01; Scale bars, 100 μm (A) and 50 μm (E).

Among the DE genes we found *Bcl6*, a transcriptional repressor that was previously reported to regulate cortical neurogenesis (Bonnefont et al., 2019; Tiberi et al., 2012), to be downregulated by 64.8% ± 0.1% in *Bcl11a* mutant neocortex (Fig. 1D). Using RNA *in situ* hybridization we found robust expression of *Bcl6* predominantly in upper and at low levels in deep cortical layers of controls between E17.5 and P4 (Fig. 1E). In *Bcl11a* mutant neocortex *Bcl6* was downregulated in upper cortical layers at these stages (Fig. 1E) suggesting this gene to be transcriptionally downstream of Bcl11a in upper cortical layers. Outside the CNS, Bcl6 exerts anti-apoptotic functions by suppressing genes involved in DNA damage response (Phan and Dalla-Favera, 2004; Phan et al., 2005; Ranuncolo et al., 2007), which could possibly be conserved in the developing neocortex as well. Therefore, we focused further analyses on Bcl6.

### Bcl6 is a direct target of Bcl11a in upper-layer cortical projection neurons

To better characterize the expression of Bcl6 protein in early postnatal somatosensory cortex we generated a polyclonal antibody in guinea pig raised against the N-terminal 484 amino acids of mouse Bcl6. Specificity of the Bcl6 antibody was tested by immunohistochemistry using *Bcl6* mutant brains, which lack exons 4-10 (Ye et al., 1997) and do not express Bcl6 protein (Tiberi et al., 2012). In comparison to wildtype littermates, we did not detect Bcl6 protein in *Bcl6* mutant brains at P0 (Fig. S3), demonstrating our antibody to specifically detect Bcl6 protein. Coexpression analysis of Bcl6 together with Bcl11a and Satb2, a marker for callosal projection neurons (Alcamo et al., 2008; Britanova et al., 2008), showed 25.6 ± 1.9% Bcl6^+^ Bcl11a^+^ Satb2^+^, 1.3 ± 0.5% Bcl6^+^ Bcl11a^+^, and 0.8 ± 0.2% Bcl6^+^ Satb2^+^ cells in wild-type brains. Only 0.3 ± 1.0% of cells exclusively expressed Bcl6 (Fig. 2A, B). Coexpression analysis of Bcl6 together with Bcl11a and Cux1, a marker for cortical layers 2-4 (Nieto et al., 2004), showed 19.0 ± 0.7% Bcl6^+^ Bcl11a^+^ Cux1^+^, 11.6 ± 0.9% Bcl6^+^ Bcl11a^+^, and 1.0 ± 0.2% Bcl6^+^ Cux1^+^ cells. Again, only 0.9 ± 0.2% of cells exclusively expressed Bcl6 (Fig. 2C, D). Thus, more than 90% of Bcl6^+^ cells coexpress Satb2 as well as Bcl11a and more than 61% of these cells are located in Cux1^+^ upper layers with distinct localization to cortical layers 2/3 (Fig. 2C). Notably, a substantial proportion of Bcl6^+^ cells is located in deep cortical layers. Thus, Bcl6 is a marker for a subset of callosal projection neurons identified by coexpression of Bcl11a and that are located in cortical layers 2/3 as well as in deep cortical layers.

**Figure 2.**
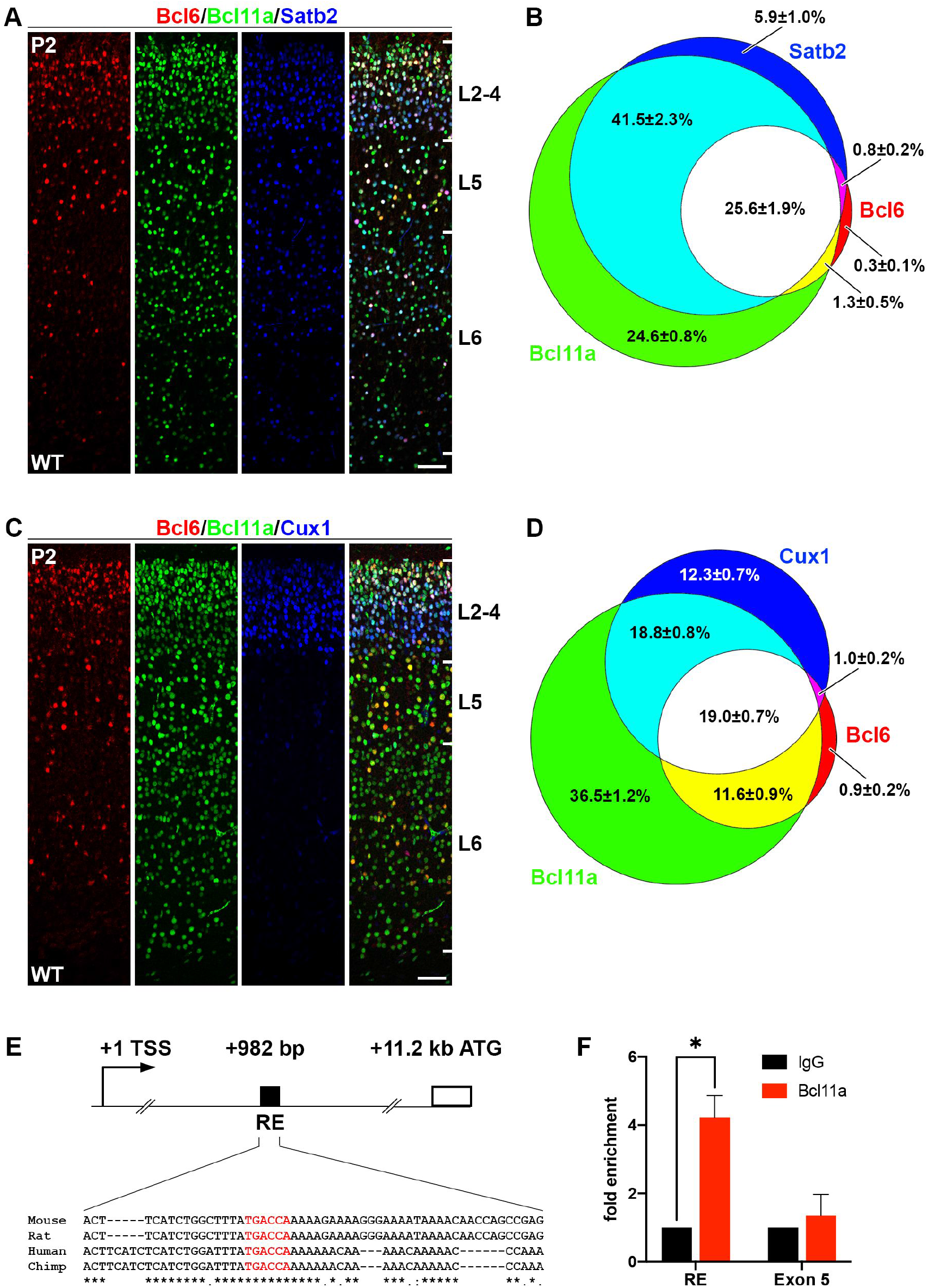
Bcl6 is expressed in superficial callosal projection neurons and a target gene of Bcl11a. (A) Immunohistochemistry of Bcl6 (red), Bcl11a (green), and Satb2 (blue) in P2 wild-type neocortex. (B) Venn diagram displaying the proportions of Bcl6 neurons overlapping with Bcl11a and Satb2 expressing cells. The percentage of each labeled cell population is given in relation to all labeled cells (Bcl6^+^ and Bcl11a^+^ and Satb2^+^, in total 4479 cells). (C) Immunohistochemistry of Bcl6 (red), Bcl11a (green), and Cux1 (blue) in P2 wild-type neocortex. (D) Venn diagram displaying the proportions of Bcl6 neurons overlapping with Bcl11a and Cux1 expressing cells. The percentage of each labeled cell population is given in relation to all labeled cells (Bcl6^+^ and Bcl11a^+^ and Cux1^+^, in total 4301 cells). (E) Scheme of the *Bcl6* gene locus displaying the start codon (ATG) at +11.2 kb relative to the transcriptional start site (TSS). A regulatory element (RE) in the first intron at +982 bp contains a conserved binding motif (TGACCA, in red) of Bcl11a. (F) ChIP analysis using a Bcl11a antibody and P2 cortical tissue detects Bcl11a binding to the RE shown in (E). Negative controls include ChIP with unspecific IgG antibody and the precipitation of exon 5 of *Bcl6* (n = 4). Student’s t test; * P < 0.05. Scale bars, 50 μm.

By DNA sequence analysis we found a TGACCA binding motif of Bcl11a (Liu et al., 2018) in the first intron that was located 982 bp downstream of the transcriptional start site and ~10.2 kb upstream of the first protein coding exon of the *Bcl6* gene. This binding motif was embedded within a 55 bp long conserved region with a high degree of conservation between rat, human, and chimp (Fig. 2E). Binding of Bcl11a to this motif was tested by chromatin immunoprecipitation (ChIP) followed by quantitative real-time PCR using a primer pair flanking this region. An enrichment of more than 4-fold was found using a Bcl11a specific antibody in comparison to an immunoglobulin G (IgG) control antibody (Fig. 2F), demonstrating binding of Bcl11a to this region. As a negative control, binding of Bcl11a to exon 5 of *Bcl6* was tested, but no significant enrichment was found in comparison to the IgG control antibody (Fig. 2F).

### Bcl6 is downregulated in upper layers of Bcl11a mutant neocortex

To confirm Bcl6 downregulation in the *Bcl11a* mutant neocortex on protein level, we performed immunohistochemistry with Bcl6 and neuron subtype specific antibodies. The overall expression of Bcl6 was reduced by 44.0% ± 7.0% compared to control neocortex at P2 (Fig. 3A, B). We did not detect changes in the number of Satb2^+^ and Cux1^+^ cells that would normally coexpress with Bcl6 (c.f. Fig 2), suggesting that these cells are born correctly, have for the most part migrated to their respective layers, and undergo neuron subtype specific differentiation (Fig. 3A, B). Furthermore, the proportion of Cux1^+^ and Satb2^+^ cells coexpressing Bcl6 was reduced from 57.4 ± 2.3% to 24.2 ± 2.6% and 58.5 ± 2.8% to 31.8 ± 1.6%, respectively, in *Bcl11a* mutant compared to control neocortex (Fig. 3A, C). As previously demonstrated, cortical thickness is reduced and layer 5 is increased at the expense of layer 6 in *Bcl11a* mutants at this stage (Wiegreffe et al., 2015; Woodworth et al., 2016). We did not detect significant changes in the number of cells coexpressing Bcl6 in deep cortical layers labelled by Bcl11b (layer 5) or Tbr1 (layer 6) (Molyneaux et al., 2007); Fig. S4) indicating a selective loss of Bcl6 in upper-layer neurons. Together, these data are compatible with a function of Bcl6 in neuron survival, which is massively impaired in upper layers of the *Bcl11a* mutant neocortex after P2 (Wiegreffe et al., 2015).

**Figure 3.**
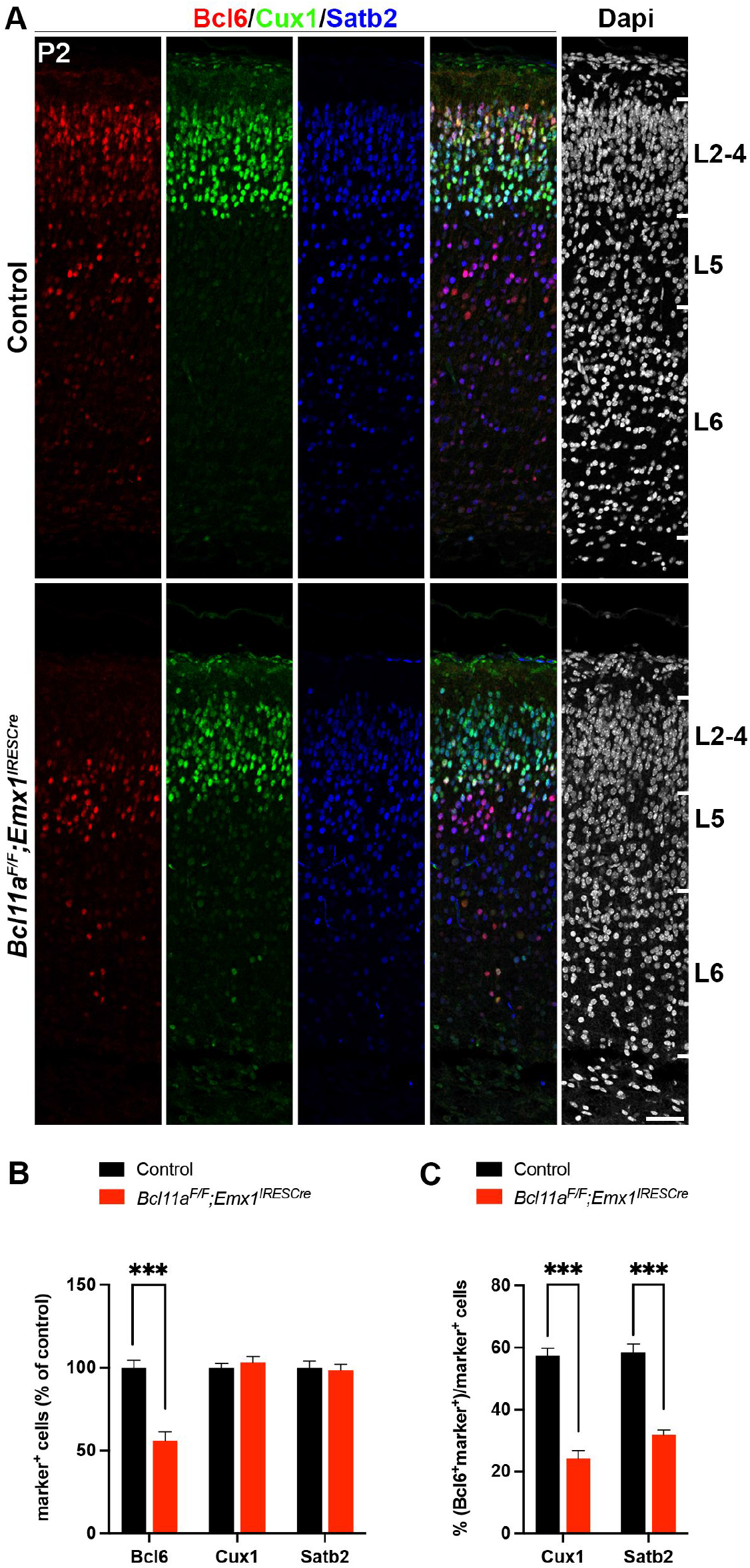
Bcl6 expression is specifically downregulated in superficial cortical layers of *Bcl11a^F/F^;Emx1^IRESCre^* neocortex. (A) Immunohistochemistry of Bcl6 (red), Cux1 (green), and Satb2 (blue) in P2 *Bcl11a^F/F^;Emx1^IRESCre^* and control neocortex. Nuclei are stained with Dapi (white). (B) Relative quantification of Bcl6^+^, Satb2^+^, and Cux1^+^ cells in *Bcl11a^F/F^;Emx1^IRESCre^* and control neocortex (n = 4). (C) Numbers of Cux1^+^ or Satb2^+^ cells that coexpress Bcl6 are reduced in *Bcl11a^F/F^;Emx1^IRESCre^* compared to control neocortex (n = 4). Student’s t test; *** p < 0.001. Scale bar, 50 μm.

### Cell-autonomous control of Bcl6 expression by *Bcl11a*

To further examine whether Bcl6 expression is directly regulated by Bcl11a in neurons, we created a mosaic mutant *in vivo* situation by using *in utero* electroporation. We generated *Bcl11a* deficient neurons in cortical layer 2/3 by electroporating Cre together with GFP (*CAG-Cre^GFP^*) or GFP alone (*CAG-Ctl^GFP^*) into conditional *Bcl11a* mutant (*Bcl11a^F/F^*) brains at E15.5 and analyzed the transfected brains at P2 (Fig. 4A, B). The proportion of GFP^+^ cells that coexpresses Bcl6 was reduced from 70.3% ± 4.0% in controls to 9.5% ± 1.5% in *Bcl11a* deficient cortical neurons (Fig. 4C, F). In contrast, the proportions of GFP^+^ cells that coexpress Cux1 or Satb2 remained unchanged (Fig. 4D-F). Thus, cell autonomous loss of *Bcl11a* in superficial cortical layers leads to a dramatic and specific reduction of Bcl6 and, together with the direct binding of Bcl11a to a conserved motif in the first intron of the *Bcl6* gene (Fig. 2E, F), suggests that Bcl11a directly controls *Bcl6* expression in these cells.

**Figure 4.**
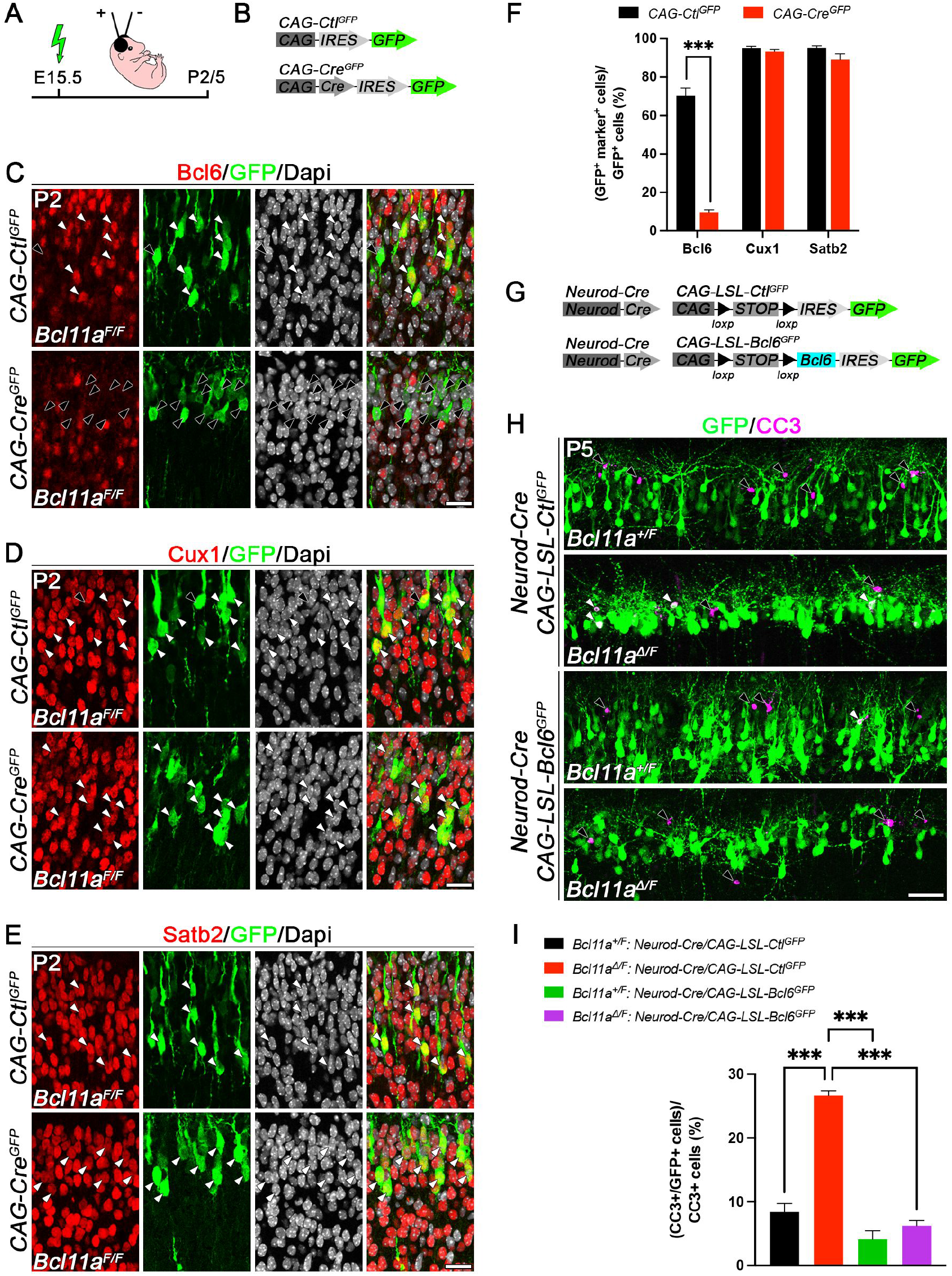
Cell autonomous loss of *Bcl11a* in superficial cortical layers leads to reduced Bcl6 expression and reintroduction of *Bcl6* into *Bcl11a* mutant superficial projection neurons rescues the *Bcl11a* mutant phenotype. (A) Schematic representation of the experimental approach. Embryos are electroporated at E15.5 with the indicated DNA plasmids and sacrificed at either P2 or P5. (B) DNA plasmids used in the experiment shown in C-F. (C-E) Immunohistochemistry of electroporated P2 *Bcl11a^F/F^* neurons in superficial cortical layers with GFP (green) and Bcl6 (red, C), Cux1 (red, D), or Satb2 (red, E) antibodies. Bcl6 expression is specifically downregulated *Bcl11a^F/F^* neocortex upon electroporation of *CAG-Cre^GFP^* in comparison to *CAG-Ctl^GFP^* control plasmid. Nuclei are stained with Dapi (white). (F) Quantification of the percentage of electroporated cells expressing Bcl6 (n = 3), Satb2 (n = 3), and Cux1 (n = 5). Results are expressed as mean ± SEM; Student’s t test; *** p < 0.001. (G) DNA plasmids used in the experiment shown in H and I. (H) Immunohistochemistry of electroporated P5 *Bcl11a^Δ/F^* and *Bcl11a^+/F^* neurons in superficial cortical layers with GFP (green) and cleaved caspase 3 (CC3, magenta) antibodies. Electroporation of *Neurod-Cre^GFP^* plasmid together with *CAG-LSL-Bcl6^GFP^* into *Bcl11a^F/F^* neocortex reduces the number of CC3^+^ cells to control levels. (I) Quantification of the experiment shown in (H) (n = 4). Results are expressed as mean ± SEM; one-way ANOVA followed by Tukey’s post hoc test; *** p < 0.001. Scale bars, 20 μm (C-E), 50 μm (H).

### Reintroduction of *Bcl6* into *Bcl11a* mutants rescues neuron death

We next asked, whether reintroduction of *Bcl6* into *Bcl11a* mutant neurons located in upper cortical layers could rescue mutant neurons from undergoing apoptosis and thereby normalize the *Bcl11a* mutant phenotype. We generated Cre-dependent control (*CAG-LSL-Ctl^GFP^*) and *Bcl6* (*CAG-LSL-Bcl6^GFP^*) expression constructs that were tested in HEK293 cells and by Western blot (Fig. 4G, S5A). Both constructs induced robust *GFP* expression in the presence or absence of *Cre*, indicating that the floxed stop (*LSL*) cassette did not prevent the *GFP* from being expressed. However, *Bcl6* expression was only observed in the presence of *Cre*, indicating a tight regulation of *Bcl6* expression from this construct (Fig. S5B). We then overexpressed *Bcl6* in *Bcl11a* mutant cortical neurons by *in utero* electroporation at E15.5 and sacrificed the brains at P5 (Fig. 4A). To circumvent functions of *Bcl6* that could interfere with neurogenesis (Bonnefont et al., 2019; Tiberi et al., 2012), we directed *Bcl6* expression to postmitotic neurons by electroporating *CAG-LSL-Ctl^GFP^* or *CAG-LSL-Bcl6^GFP^* expression constructs together with *Cre* placed under the control of the postmitotically activated *Neurod* promoter (*Neurod-Cre*) into *Bcl11a^Δ/F^* (i.e. conditional mutant) or *Bcl11a^+/F^* (i.e. control) brains (Fig. 4G). Co-electroporation of *Neurod-Cre* together with *CAG-LSL-Ctl^GFP^* robustly induced cell death in *Bcl11a^Δ/F^* in comparison to control brains by more than 3-fold. In contrast, postmitotic reintroduction of *Bcl6* into *Bcl11a* deficient neurons reduced apoptosis to control levels. Of note, overexpression of *Bcl6* in control brains did not significantly reduce the number of cleaved caspase 3^+^ cells below control levels (Fig. 4H, I). Together, these data strongly support a role for *Bcl6* as a direct functional downstream target of Bcl11a that plays an important role for neuron survival during the second wave of developmental cell death at the early postnatal stage.

### Increased cell death in postnatal *Bcl6* mutant neocortex

To further corroborate that *Bcl6* confers survival of cortical projection neurons, we generated forebrain-specific *Bcl6* mutants by crossing conditional *Bcl6* mutant mice (*Bcl6^F/F^*), in which exons 7-9 are flanked by *loxP* sites (Hollister et al., 2013), with *Nex^Cre^* mice (Goebbels et al., 2006) that induce recombination in postmitotic cortical projection neurons. Quantitative real-time PCR showed that *Nex^Cre^* reduced *Bcl6* expression by 80.0% ± 0.1% compared to controls at P0 (Fig. 5B). Due to restricted activity of *Nex^Cre^* incomplete reduction of Bcl6 is most likely caused by residual expression in non-neuronal cell types. We chose P5 to analyze developmental cell death in *Bcl6* mutant brains because *Bcl11a* mutants display massively increased cell death (Wiegreffe et al., 2015) and naturally occurring cell death in wildtype brains peaks around this stage (Blanquie et al., 2017). We found a significant increase of cleaved caspase 3^+^ cells located predominantly in the upper cortical layers from 6.04% ± 0.02% in controls to 8.44% ± 0.77% cells/mm^2^ in *Bcl6* mutant brains concomitant with a reduction of cortical area by 9.7% ± 2.0% (Fig. 5A, C, D). Collectively, these data show that Bcl6 exerts functions in upper-layer neuron survival during early postnatal neocortical development. To characterize downstream genetic pathways of Bcl6 responsible for the observed phenotype, we isolated upper cortical layers from *Bcl6* mutant and control brains by laser capture microdissection at P5 and performed a differential expression analysis on this tissue using microarrays. This analysis revealed a small number of DE genes that were mostly upregulated in *Bcl6* mutant upper cortical layers (Fig. 5E). Of these genes, the cell death associated factor Foxo1, previously demonstrated to be involved in regulation of neuron death (Carter and Brunet, 2007; Santo and Paik, 2018) was found upregulated in *Bcl6* as well as in *Bcl11a* mutants. Differential expression was verified by quantitative real-time PCR and RNA *in situ* hybridization. *Foxo1* expression was upregulated more than 2-fold (Fig. 5F) and upregulation was most apparent in upper layers of the *Bcl6* mutant neocortex compatible with an induction of cell death at this stage. In lymphoid cells Bcl6 regulates cell death through p53 function (Cerchietti et al., 2008; Phan and Dalla-Favera, 2004). We did not detect changes in p53 expression in our expression analysis.

**Figure 5.**
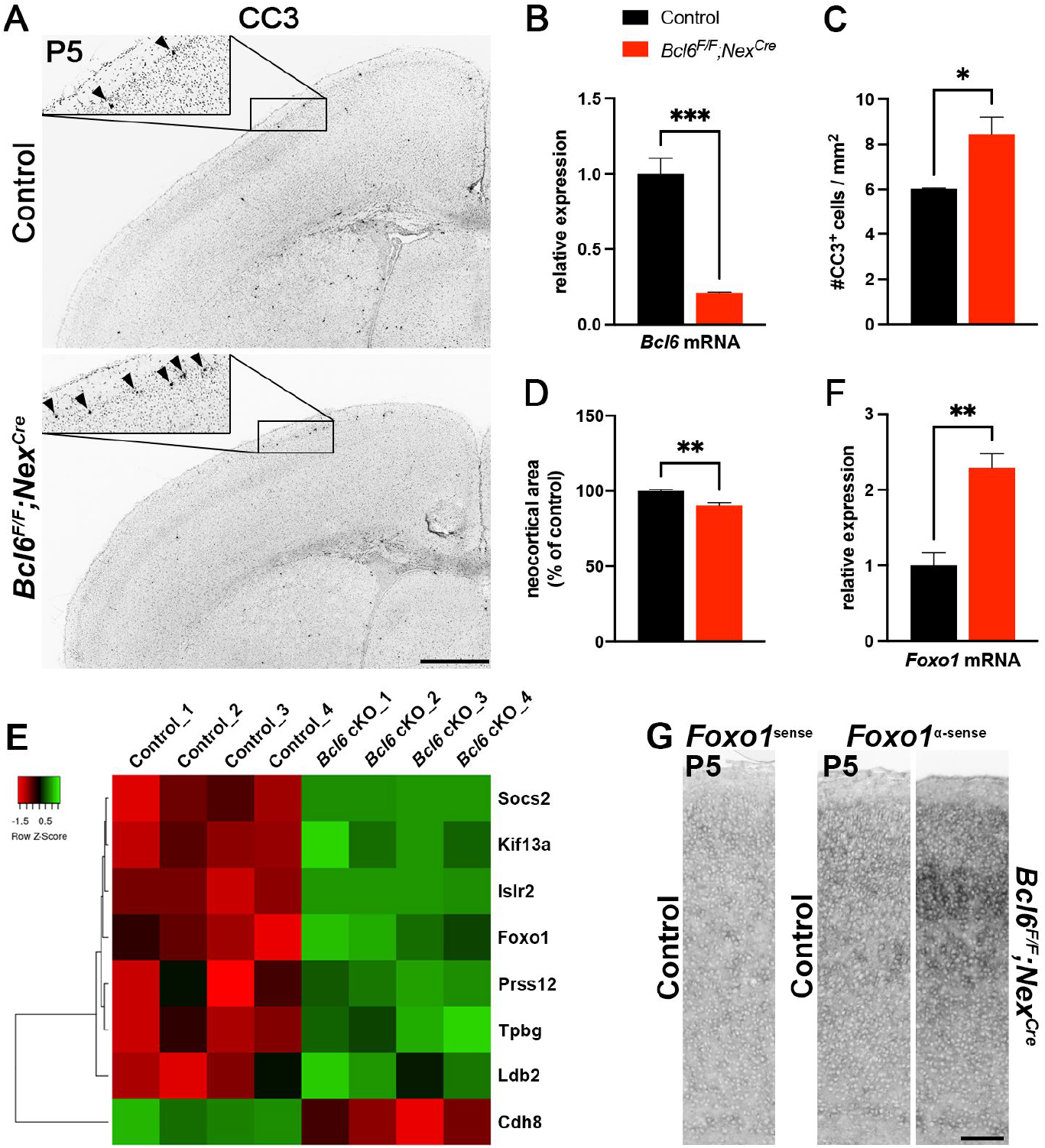
Postnatal developmental cell death is increased in *Bcl6^F/F^;Nex^Cre^* neocortex. (A) Immunohistochemistry of cleaved caspase 3 (CC3) shows that the number of CC3^+^ cells is increased in P5 *Bcl6^F/F^;Nex^Cre^* compared to control neocortex. Insets are enlargements of the boxed areas in corresponding panels. (B) Relative *Bcl6* mRNA expression level determined by quantitative real-time PCR using primers targeting a region of exon 8 is decreased in E15.5 *Bcl6^F/F^;Nex^Cre^* compared to control neocortex. (C) Quantification of the experiment shown in (A) (n = 3). (D) Quantification of neocortical area in P5 *Bcl6^F/F^;Nex^Cre^* and control brains (n = 3). (E) Heat map showing differentially expressed genes in laser microdissected superficial cortical layers of P5 *Bcl6^F/F^;Nex^Cre^* compared to control brains (n = 4). (F) Relative *Foxo1* mRNA expression level determined by quantitative real-time PCR is increased in laser microdissected superficial cortical layers of P5 *Bcl6^F/F^;Nex^Cre^* compared to control brains (n = 4). (G) RNA *in situ* hybridization showing upregulation of *Foxo1* expression in P5 *Bcl6^F/F^;Nex^Cre^* compared to control neocortex. All graphs represent mean ± SEM; Student’s t test; * p > 0.05; ** p < 0.01. Scale bars, 500 μm (A), 50 μm (G).

Together, our data uncover *Bcl6* as a direct functional downstream target of Bcl11a that plays an important role in regulation of DCD of upper-layer CPN at early postnatal stages.

## Discussion

Selective deletion of the zinc finger transcription factor Bcl11a in CPN results in massively increased developmental neuron death during early postnatal development of the murine neocortex (Wiegreffe et al., 2015). In the present study we used this genetic model as a tool to detect candidate regulatory mechanisms in control of DCD during cortical development. We identified *Bcl6* as a direct downstream transcriptional target of Bcl11a. Bcl6 is downregulated in *Bcl11a* mutant upper-layer CPN. Reintroduction of *Bcl6* into *Bcl11a* mutant upper-layer CPN completely rescues the apoptosis phenotype, while conditional deletion of *Bcl6* in CPN induces neuron death. Finally, our data suggest Bcl6 to exert part of its survival functions through *Foxo1*. Together, in the present study we uncover a novel, neuron-type and developmental-phase specific Bcl11a/Bcl6-dependent transcriptional pathway in the control of developmental neuron death in murine neocorticogenesis.

During neocorticogenesis DCD occurs in two spatiotemporally defined waves. In mice, numbers of CPN have been shown to be refined by DCD between P4 and P6 (Blanquie et al., 2017; Verney et al., 2000; Wong et al., 2018). This time course precisely corresponds to elevated apoptosis of upper-layer CPN observed in *Bcl11a* mutants (Wiegreffe et al., 2015), demonstrating our genetic approach to be phase- and cell-type specific. Comparative transcriptomic analyses revealed 137 differentially expressed genes. Among these candidates, the BTB/POZ zinc finger transcriptional regulator Bcl6 was found to be downregulated in mutants. This phenotype was restricted to Cux1 and Satb2 expressing CPN within upper cortical layers 2 and 3. Further analysis revealed that Bcl11a directly controls *Bcl6* expression through binding to a conserved element within the *Bcl6* promotor.

Previously, we demonstrated Bcl6 to be required during early phases of neocortical, where Bcl6 promotes the transition of neural progenitors into postmitotic neurons (Bonnefont et al., 2019; Tiberi et al., 2012). Our data suggest additional functions of Bcl6 in the postnatal development of postmitotic CPN. A conserved function of this factor in control of cell survival is supported by its well-characterized functions in the lymphatic system. Bcl6 prevents apoptosis in germinal center B-cells and exerts oncogenic activity in diffuse large B-cell lymphoma both, through modulation of the p53 downstream pathway (Cerchietti et al., 2008; Phan and Dalla-Favera, 2004). In the cerebellum, deletion of *Bcl6* induces massively increased cell death of granule cell precursors but not postmitotic granule cells leading to reduction of organ size (Tiberi et al., 2014). Interestingly, activation of nuclear calcium pathway through synaptic NMDA-receptor signaling induces Bcl6 expression in hippocampal neurons. In turn, upregulation of Bcl6 improves survival of these neurons (Zhang et al., 2007). This suggests, that activity-dependent as well as activity-independent, transcriptional regulatory pathways converge onto Bcl6 in the control of DCD.

Compared to *Bcl11a* mutants we observed only moderate increase in apoptosis in *Bcl6* mutant CPN, raising the possibility of additional signals to contribute to apoptosis in *Bcl11a* mutants. Postnatally, *Bcl11a* mutant CPN display severe morphogenetic defects as characterized by shortened apical dendrites and disturbed dendritic branching pattern (Wiegreffe et al., 2015). This may result in impaired synaptic integration and electrical activity of *Bc11a* mutant neurons and in turn contribute to the severity of the phenotype. Alternatively, additional, not yet characterized signals, might be involved.

In our screen, we detected several axon guidance molecules, including Slit2, Efna5, Sema3c, -3d, -7a, Flrt2, -3 to be deregulated in *Bcl11a* mutant CPN. Semaphorins, for example, have extensively been demonstrated to influence neuronal connectivity (Koropouli and Kolodkin, 2014). Thus, differentially expressed guidance molecules, as observed in our study, might either directly or indirectly, through modulation of connectivity influence the severity of the apoptosis phenotype in *Bcl11a* mutants. In addition, we found cadherin 6, 12, 13 and protocadherin 9 to be deregulated in *Bcl11a* mutant CPN. Recent experimental evidence suggests cadherins, in addition to their well characterized functions in cell recognition and neural circuit assembly (Jontes, 2018; Sanes and Zipursky, 2020) to exert survival functions, for example in neocortical interneurons as well (Lazlo 2020).

In lymphoid cells Bcl6 regulates cell survival through modulation of the p53 pathway (Cerchietti et al., 2008; Phan and Dalla-Favera, 2004). While p53 expression was found unchanged in both, *Bcl11a* and in *Bcl6* mutant CPN, significant upregulation of *Foxo1* in both *Bcl6* and *Bcl11a* mutants was detected. Bcl11a and Bcl6 were shown to physically interact and colocalize in nuclear paraspeckles suggesting common regulation of gene expression (Liu et al., 2006; Nakamura et al., 2000). Members of the Foxo family have been demonstrated to be involved in the control of neuron survival (Carter and Brunet, 2007; Santo and Paik, 2018). It might thus well be, that Bcl6 together with Bcl11a exerts anti-apoptotic functions in CPN through this pathway.

The extent of DCD is specific to individual cortical areas. For example, numbers of neurons eliminated by DCD are substantially higher in the somatomotor as compared to the somatosensory cortex (Blanquie et al., 2017). Loss of *Bcl11a* in the neocortex has been previously suggested to result in changed area identity, characterized by the partial motorization of the *Bcl11a* mutant neocortex defined on a transcriptional level (Greig et al., 2016). Intriguingly, Bcl6 expression in the wildtype neocortex at P4 is lower in the somatomotor as in the somatosensory cortex (Greig et al., 2016). Thus, the identification of a Bcl11a/Bcl6-dependent regulatory pathway of DCD in this study provides additional mechanistic insights into how Bcl11a contributes to establishing cortical area identity.

## Experimental Procedures

### Animals

Mice carrying a conditional knockout allele of *Bcl11a* (*Bcl11a^F^*) have previously been described (John et al., 2012). These mice were crossed to *Emx1^IRESCre^* or *Deleter^Cre^* (Gorski et al., 2002; Schwenk et al., 1995) mice to generate *Bcl11a^Δ/+^* heterozygous and conditional *Bcl11a^F/F^;Emx1^IRESCre^* mutants, respectively. *Bcl11a^F/+^;Emx1^IRESCre^* littermates served as controls. Mice carrying a conditional knockout allele of *Bcl6* (*Bcl6^F^*) were crossed to *Nex^Cre^* mice (Goebbels et al., 2006; Hollister et al., 2013) to generate conditional *Bcl6^F/F^;Nex^Cre^* mutants. *Bcl6^F/F^* littermates without a *Nex^Cre^* allele served as controls. *Bcl6^+/−^* mice have previously been described (Ye et al., 1997). Genotyping of the mice was performed by PCR. All mouse experiments were carried out in compliance with German law and approved by the respective government offices in Tübingen, Germany.

### Immunohistochemistry and RNA *in situ* hybridization

Brains were fixed in 4% PFA in 0.1M phosphate buffer (pH 7.4), embedded in OCT compound (Polysciences), and frozen sections were prepared at 14 μm for immunohistochemistry or 18 μm for RNA *in situ* hybridization as previously described (John et al., 2012; Simon et al., 2012). Paraffin and vibratome sections were prepared at 7 μm and 50 μm, respectively. All clones for non-radioactive RNA *in situ* hybridization, except for Flrt2 and Flrt3, which were a gift by Rüdiger Klein (Max Planck Institute of Neurobiology, Martinsried, Germany), were generated by reverse transcription PCR and oligonucleotides are listed in supplemental table 1.

The following antibodies were used: guinea pig anti-Bcl11a (John et al., 2012), mouse anti-Bcl11a (Abcam Cat# ab19487, RRID:AB_444947), rabbit anti-Bcl11a (John et al., 2012), rat anti-Bcl11b (Abcam Cat# ab18465, RRID:AB_2064130), goat anti-Brn2 (Santa Cruz Biotechnology Cat# sc-6029, RRID:AB_2167385), rabbit anti-cleaved Caspase 3 (Cell Signaling Technology Cat# 9661, RRID:AB_2341188), rabbit anti-Cux1 (Santa Cruz Biotechnology Cat# sc-13024, RRID:AB_2261231), chicken anti-GFP (Abcam Cat# ab13970, RRID:AB_300798), mouse anti-Satb2 (Abcam Cat# ab51502, RRID:AB_882455), and rabbit anti-Tbr1 (Abcam Cat# ab31940, RRID:AB_2200219). To generate anti-Bcl6 antiserum guinea pigs were injected with a protein comprising amino acids 4-484 of mouse Bcl6 (NP_033874) and pooled sera were purified by affinity chromatography. Biotin-conjugated, HRP-conjugated, and fluorescent secondary antibodies were purchased from Jackson ImmunoResearch. Sections were counterstained with Dapi (Molecular Probes). Immunohistochemical detection of Bcl6 was performed on paraffin sections with antigen retrieval by boiling the section for 30 minutes in Tris-EDTA buffer, pH 9.0 and enhanced using tyramide signal amplification (Invitrogen) according to the manufacturer’s instructions or an avidin/biotin-based peroxidase system and DAB substrate (Vector Laboratories). Cleaved caspase 3 was detected on frozen sections of conditional *Bcl6* mutants using an avidin/biotin-based peroxidase system and DAB substrate (Vector Laboratories). All fluorescent images were examined on a TCS SP5II confocal microscope (Leica) and processed with Adobe Photoshop (RRID:SCR_014199) software.

### Laser microdissection

All procedures were performed in an RNase-free environment. Cortical layers 2-4 were isolated from unfixed frozen sections via laser microdissection. Briefly, brains were quickly removed from the skull, washed in ice-cold PBS, frozen in OCT compound (Polysciences), and stored at −80°C. Sections were prepared at 20 μm and mounted on membrane-covered 1 mm PEN slides (Zeiss) that were UV-treated and coated with poly-L-lysine. Sections were fixed in ice-cold 70% EtOH for 1 min, incubated in 1% cresyl violet acetate solution (Waldeck) for 45 sec, and washed in 70% EtOH and 100% EtOH for 1 min each. After a brief drying step on a 37°C warming plate, sections were immediately processed for laser microdissection using a PALM MicroBeam Rel.4.2 (Zeiss). Laser microdissected tissue was lysed in RLT lysis buffer (Qiagen) containing 2-mercaptoethanol for 30 min on ice and stored at −80°C before total RNA extraction.

### Plasmids

*CAG-Ctl^GFP^* and *CAG-Cre^GFP^* have previously been described (Hand et al., 2005; Wiegreffe et al., 2015). The recombinase *Cre* was from *CAG-Cre^GFP^* and inserted into *pNeuroD-ires-GFP* (gift of Franck Polleux; RRID:Addgene_61403) to generate *NeuroD-Cre*. The *ires-GFP* cassette was cut from *CAG-Ctl^GFP^* and inserted into *pCALNL-GFP* (gift of Connie Cepko; RRID:Addgene_13770). *Bcl6* (NM_009744) was cloned by PCR using a cDNA clone (Cat.No. MC203091, Origene) as template and inserted into *CAG-Ctl^GFP^* to generate *CAG-LSL-Bcl6^GFP^*. *CAG-Cre* was a gift of Connie Cepko (RRID:Addgene_61403).

### *In utero* electroporation

*In utero* electroporation was performed as previously described (Saito and Nakatsuji, 2001; Wiegreffe et al., 2017) with minor modifications. Briefly, pregnant dams were anaesthetized with Isoflurane (Abbott) and 1-2 μL of plasmid DNA were injected per embryo at a concentration of 0.5-1.0 μg/μL per construct. 5 mm electrodes (Nepagene) and five pulses of 40 V (50 ms ON, 950 ms OFF) generated by a CUY21 EDIT electroporator (Nepagene) were used to transfect cells in the dorsolateral ventricular zone.

### Microarray analysis, quantitative real-time PCR, and chromatin immunoprecipitation

Microarray analysis was performed as previously described (John et al., 2012; Simon et al., 2012) with minor modifications. Briefly, total RNA was isolated from laser-microdissected control and mutant samples (n = 4) using the RNeasy Micro Plus Kit (Qiagen). The isolated RNA was checked for purity and integrity using Nanodrop spectrophotometer and TapeStation (Agilent), respectively. Transcriptome analysis was performed using GeneChip Mouse Gene 1.0 ST Arrays (Affymetrix) and BRB-ArrayTools developed by Dr. Richard Simon and BRB-ArrayTools Development Team (http://linus.nci.nih.gov/BRB-ArrayTools.html). The data obtained in our microarray experiments were deposited at the GEO website under accession numbers GEO: GSE185287 and GSE185288.

Total RNA was reverse transcribed using the SensiFast cDNA Synthesis Kit (Bioline), and quantitative real-time PCR was performed using the LightCycler DNA Master SYBR Green I Kit in a LightCycler 480 System (Roche). Oligonucleotides used for quantitative real-time OCR are listed in supplemental table S2. The relative copy number of *Gapdh* RNA was quantified and used for normalization. Data were analyzed using the comparative CT method (Schmittgen and Livak, 2008).

Chromatin immunoprecipitation (ChIP) was carried out as previously described (Nelson et al., 2006) with minor modifications. Briefly, P0 cortical tissue was collected from wild-type pups, flash frozen in liquid nitrogen, and stored at −80°C until ChIP. Tissue was disrupted in low sucrose buffer (320mM sucrose, 10mM HEPES, pH 8.0, 5mM CaCl2, 3mM Mg[CH3COO]2, 1mM DTT, 0.1mM EDTA, 0.1% Triton X-100) and fixed for 15 min at RT in 1% formaldehyde. After quenching with glycine solution, nuclei were washed in Nelson buffer (140mM NaCl, 20mM EDTA, pH 8.0, 50mM Tris, pH 8.0, 1% Triton X-100, 0.5% NP-40) and disrupted in RIPA buffer (140 mM NaCl, 10mM Tris, pH 8.0, 1mM EDTA, pH 8.0, 1% SDS, 1% Triton X-100, 0.1% NaDOC). Chromatin was sonicated for 40 cylces (30 sec ON/OFF) using a Bioruptor Plus (Diagenode) with high power settings. For each ChIP reaction 15 μg of sheared chromatin was diluted ten times with IP buffer (50mM Tris, pH 8.0, 150 mM NaCl, 1% NP-40, 0.5% NaDOC, 20mM EDTA, pH 8.0, 0.1% SDS) and incubated overnight at 4°C with 3 μL specific mouse monoclonal antibody recognizing Bcl11a (Abcam Cat# ab19487, RRID:AB_444947) or unspecific IgG1 antibody (Cell Signaling Technology Cat# 5415, RRID:AB_10829607), which served as a negative control. 20 μL of protein G magnetic beads (Invitrogen) were added to each ChIP reaction for two hours at 4°C. After washing with IP buffer containing 0.1% SDS, LiCl buffer (500mM LiCl, 100mM Tris, pH 8.0, 1% NP-40, 1% NaDOC, 20mM EDTA, pH 8.0), and TE buffer (10mM Tris, pH 8.0, 1mM EDTA, pH 8.0), DNA was eluted from beads and purified by phenol-chloroform extraction. The precipitated DNA was analyzed by quantitative real-time PCR using oligonucleotides recognizing a conserved Bcl11a binding motif (TGACCA) in the first intron of *Bcl6*. As negative controls, oligonucleotides were used recognizing a region of exon 5 of *Bcl6* and the *Hprt* promoter region, respectively. All oligonucleotide sequences are listed in supplemental table S1. ChIP quantitative real-time PCR data were analyzed by the comparative CT method determining the fold enrichment of the immunoprecipitated DNA by the specific antibody versus IgG1 using the input as a reference.

### Cell culture and western blotting

HEK293 cells were grown in DMEM with 10% fetal calf serum and 1% penicillin/streptomycin at 37°C under 5% CO_2_ atmosphere. Cells were transfected using Lipofectamine 2000 according to the manufacturer’s instructions (Invitrogen). Total proteins were extracted with ice-cold lysis buffer (1% NP-40, 150mM NaCl, 50mM Tris, pH 8.0, 1mM EDTA), separated by SDS-PAGE, and electrophoretically transferred onto PVDF membranes (Amersham). Membranes were blocked with 5% non-fat milk (Bio-Rad) and incubated with mouse anti-beta-actin (Abcam Cat# ab8226, RRID:AB_306371), rabbit anti-Bcl6 (Santa Cruz Biotechnology Cat# sc-858, RRID:AB_2063450), and chicken anti-GFP (Abcam Cat# ab13970, RRID:AB_300798), followed by treatment with horseradish peroxidase-conjugated secondary antibodies (Jackson ImmunoResearch) and ECL Plus western blotting detection reagents according to the manufacturer’s instructions (ThermoScientific).

### Cell counts and statistical analysis

For each experiment, at least three control and three mutant brains were analyzed, and three to five sections per brain were quantified. Anatomically matched sections were selected from an anterio-posterior level between the anterior commissure and the dorsal hippocampus. Stained cells were counted in radial units of 100 μm (Fig. 2, 3, S5), 350 μm (Fig. 4C-E), or 750 μm (Fig. 4H) width in the presumptive somatosensory cortex or in the entire neocortex (Fig. 5). Cells were counted using ImageJ (RRID:SCR_003070) and Imaris (RRID:SCR:007370) software. Statistical analysis was done with Microsoft Excel (RRID:SCR_016137) or GraphPad Prism (RRID:SCR_002798) software. Venn diagrams were generated using MATLAB (RRID:SCR_001622) software. Significance between groups was assessed using a two-tailed Student’s t test or one-way ANOVA, followed by Tuckey’s post hoc test. P values < 0.05 were considered statistically significant.

## Acknowledgements

We are grateful to C. Cepko (Harvard Medical School, Boston), K.-A. Nave (Max-Planck-Institute for Experimental Medicine, Göttingen), F. Polleux (Columbia University, New York) for the gift of mice and providing DNA plasmids. We thank K. Holzmann of the core facility “Genomics” and the staff of core facility “Laser Microdissection” of the Medical Faculty of Ulm University. We thank J. Andratschke, L. Schmid, and D. Krattenmacher for excellent technical assistance. This work was supported by grants from the Deutsche Forschungsgemeinschaft to S.B. (BR 2215/1-2), Ulm University (Bausteinprogramm 3.2) to C.W. and the German Academic Schlolarship Foundation to T.W.

## Author contributions

C.W. and S.B. designed the experiments. C.W., T.W., N.J., conducted the experiments and analyzed the data. J.B. and P.V. provided mice and reagents. C.W. and S.B. wrote the manuscript

## Supplementary figure legends

**Supplementary Figure S1.**
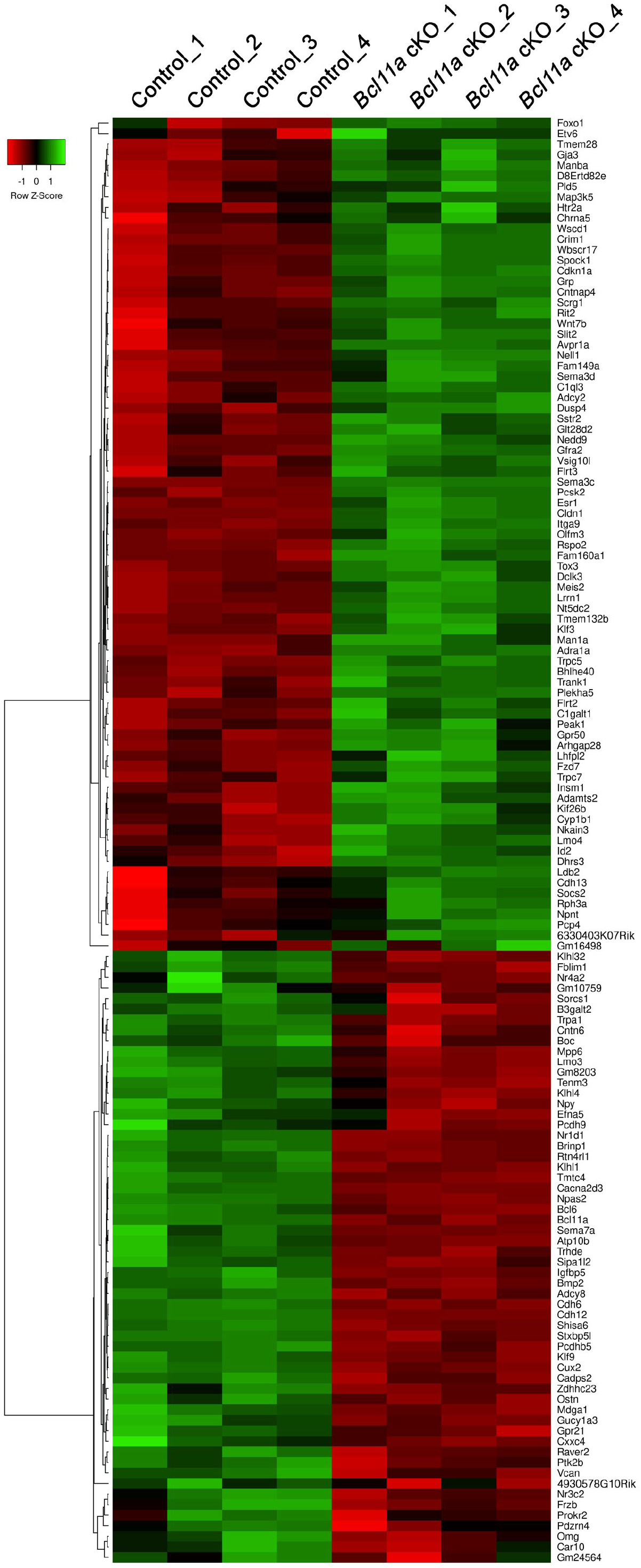
Heat map of differentially expressed genes in upper layers of P2 *Bcl11a^F/F^;Emx1^IRESCre^* compared to control neocortex. Heat map showing 79 upregulated genes in green and 58 downregulated genes in red in upper neocortical layers conditional *Emx1^IRESCre^;Bcl11a^F/F^* mutants (*Bcl11a* cKO) compared to controls at postnatal day 2.

**Supplementary Figure S2.**
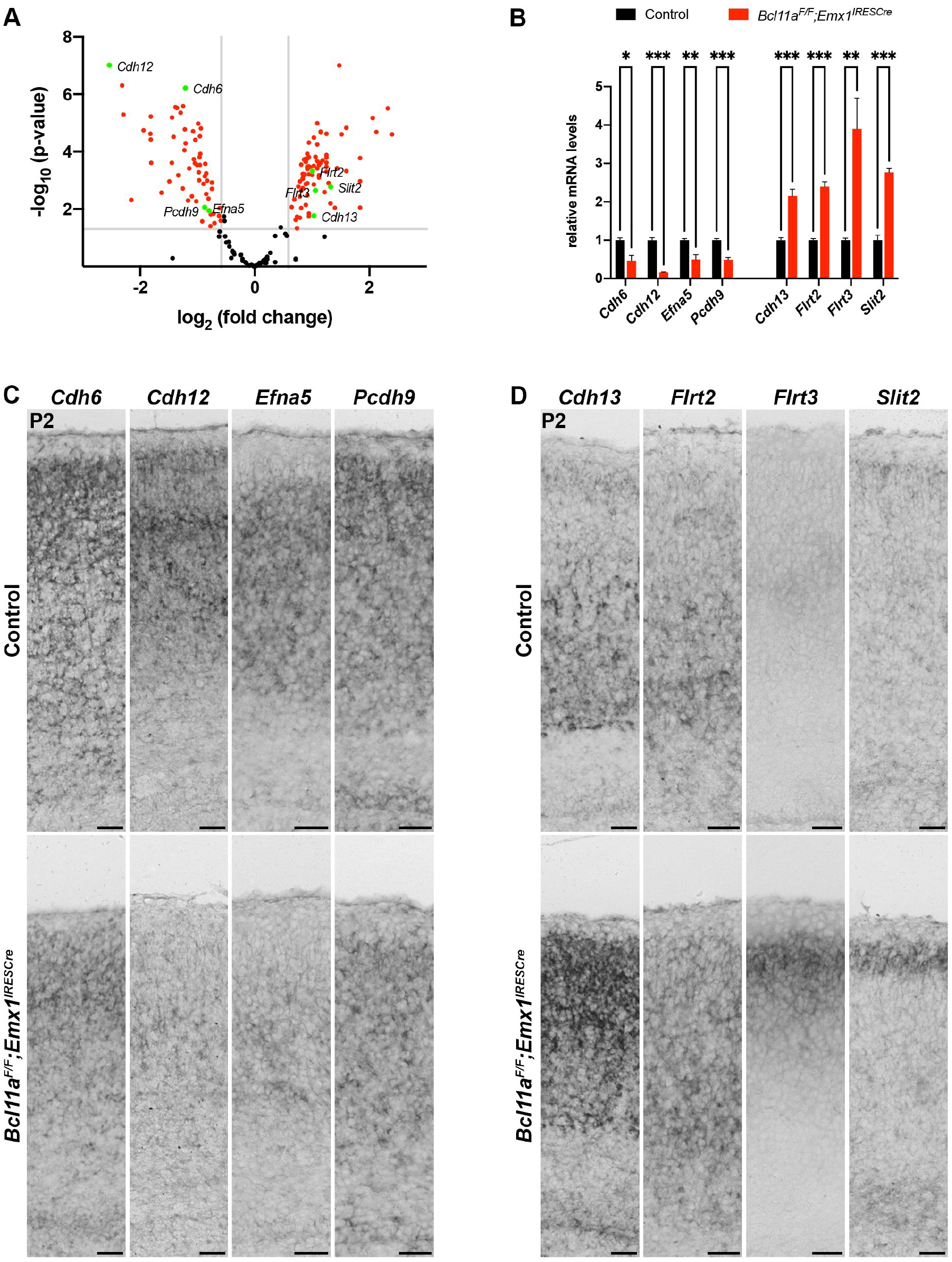
Selected candidate downstream target genes of Bcl11a in superficial cortical layers at early postnatal development. (A) Volcano plot showing differentially expressed (DE) genes in laser microdissected cortical layers 2-4 of *Bcl11a^F/F^;Emx1^IRESCre^* neocortex compared to controls. DE genes not significantly changed (fold change < 1.5; *p* > 0.05) are shaded black. *Cdh6*, *Cdh12*, *Efna5*, *Pcdh9*, *Cdh13*, *Flrt2*, *Flrt3*, and *Slit2* are highlighted in green. (B) Relative mRNA expression levels of select DE genes determined by quantitative real-time PCR in laser microdissected cortical tissue of P2 *Bcl11a^F/F^;Emx1^IRESCre^* and control brains (n = 4). Graph represents mean ± SEM; Student’s t test; * p > 0.05; ** p < 0.01; *** p < 0.01. (C) RNA *in situ* hybridization of selected DE genes with decreased expression in P2 *Bcl11a^F/F^;Emx1^IRESCre^* compared to control neocortex. (D) RNA *in situ* hybridization of select DE genes with increased expression in P2 *Bcl11a^F/F^;Emx1^IRESCre^* compared to control neocortex. Scale bar, 50 μm.

**Supplementary Figure S3.**
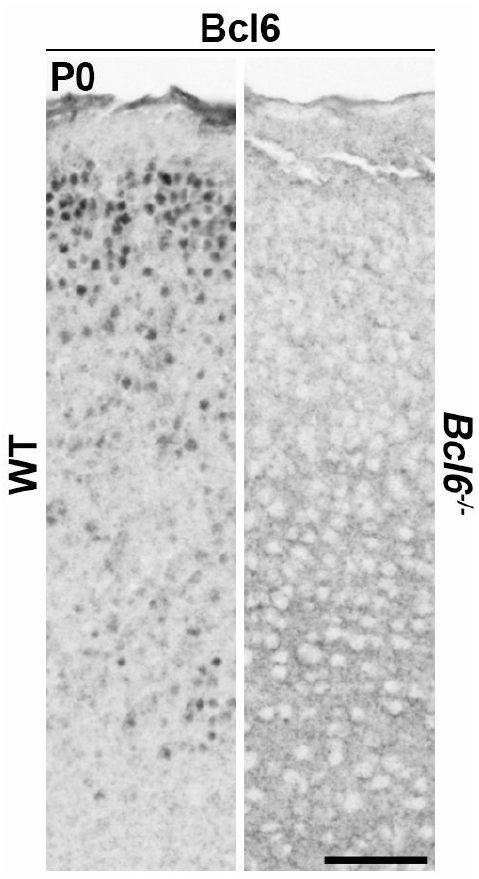
Specificity of Bcl6 antibody shown by immunohistochemistry. Immunohistochemistry of Bcl6 using guinea pig anti-Bcl6 antibody in P0 *Bcl6^−/−^* and wildtype neocortex. Bcl6 epxression is detectable in upper layers and at lower levels in deep layers of wildtype neocortex. In *Bcl6^−/−^* mutants Bcl6 expression is not detectable. Scale bar, 50 μm.

**Supplementary Figure S4.**
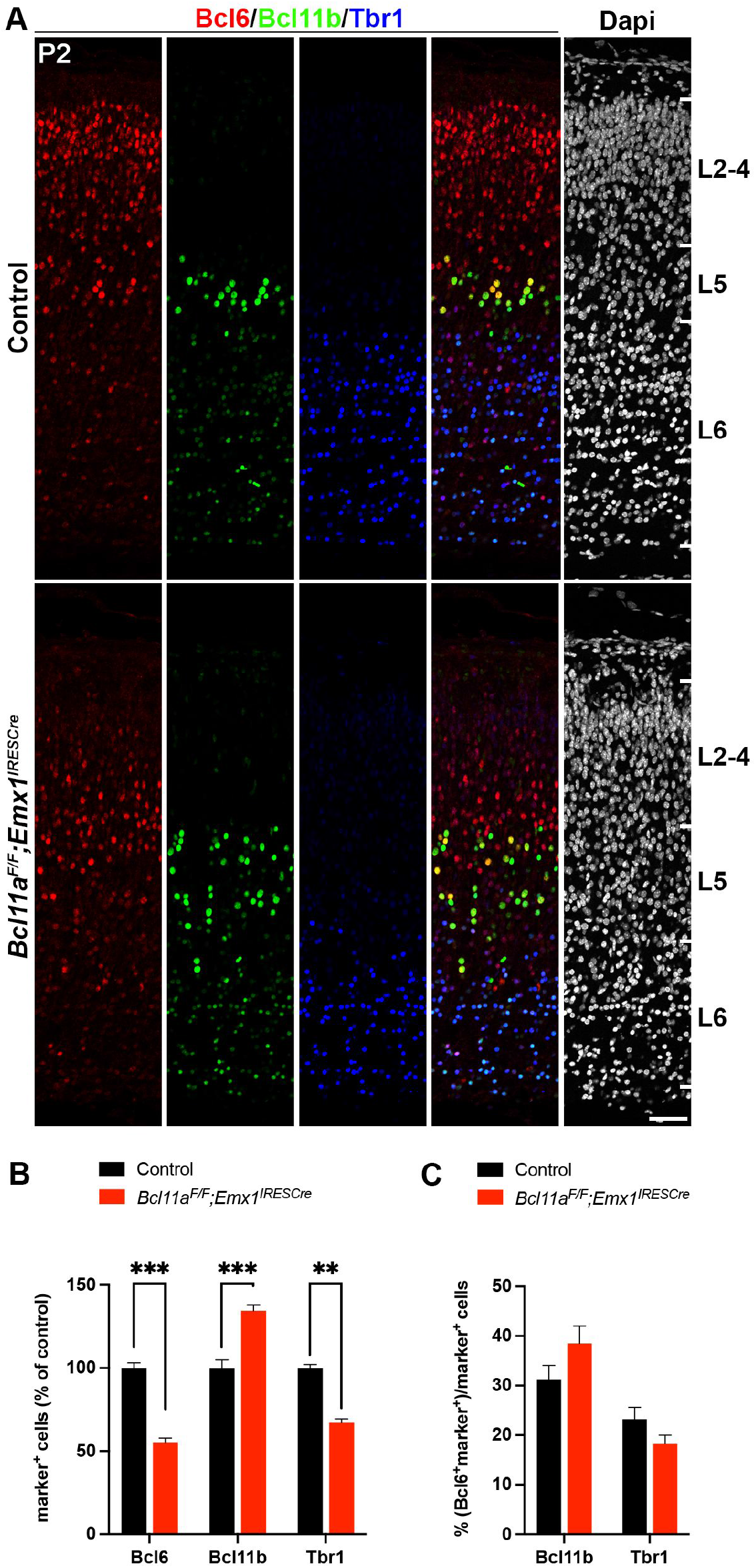
Relative Bcl6 expression is unchanged in deep cortical layers of *Bcl11a^F/F^;Emx1^IRESCre^* compared to control neocortex. (A) Immunohistochemistry of Bcl6 (red), Bcl11b (green), and Tbr1 (blue) in P2 *Bcl11a^F/F^;Emx1^IRESCre^* and control neocortex. Bcl6 and Tbr1 expressions are downregulated while Bcl11b expression is upregulated in *Bcl11a^F/F^;Emx1^IRESCre^* compared to control neocortex. Nuclei are stained with Dapi (white). (B) Relative quantification of Bcl6^+^, Bcl11b^+^, and Tbr1^+^ cells in *Bcl11a^F/F^;Emx1^IRESCre^* and control neocortex (n = 4). (C) Numbers of Bcl11b^+^ or Tbr1^+^ cells that coexpress Bcl6 are normal in *Bcl11a^F/F^;Emx1^IRESCre^* compared to control neocortex (n = 4). Student’s t test; ** p < 0.01; *** p < 0.001. Scale bar, 50 μm.

**Supplementary Figure S5.**
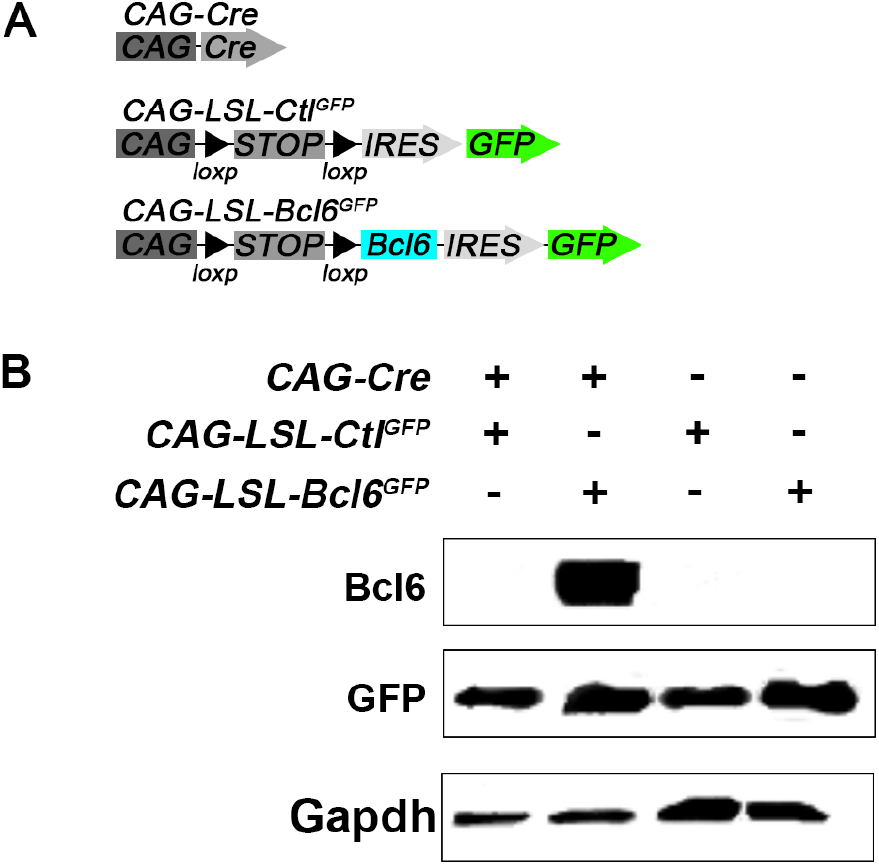
Western blot analysis of Cre-dependent DNA plasmids for overexpression of Bcl6 cDNA. (A) DNA plasmids used for transfection of HEK293 cells. (B) Western Blot analysis of total protein lysates shows that *CAG-LSL-Bcl6^GFP^* plasmid expresses *Bcl6* only in the presence *CAG-Cre* plasmid. Note that GFP is expressed independent of *pCAG-Cre*. Gapdh served as protein loading control.

**Supplementary Table 1: Differentially expressed gene in upper cortical layers of P2 *Bcl11a* cKO compared to controls**

List of the genes identified by microarray analysis that are differentially expressed in upper layers of P2 *Emx1^IRESCre^;Bcl11a^F/F^* (*Bcl11a* cKO) compared to control neocortex.

**Supplementary Table 2: Primers used in this study.**

